# Measuring the distribution of fitness effects in somatic evolution by combining clonal dynamics with dN/dS ratios

**DOI:** 10.1101/661264

**Authors:** Marc J Williams, Luiz Zapata, Benjamin Werner, Chris Barnes, Andrea Sottoriva, Trevor A Graham

## Abstract

The distribution of fitness effects (DFE) defines how new mutations spread through an evolving population. The ratio of non-synonymous to synonymous mutations (dN/dS) has become a popular method to detect selection in somatic cells, however the link, in somatic evolution, between dN/dS values and fitness coefficients is missing. Here we present a quantitative model of somatic evolutionary dynamics that yields the selective coefficients from individual driver mutations from dN/dS estimates, and then measure the DFE for somatic mutant clones in ostensibly normal oesophagus and skin. We reveal a broad distribution of fitness effects, with the largest fitness increases found for TP53 and NOTCH1 mutants (proliferative bias 1-5%). Accurate measurement of the per-gene DFE in cancer evolution is precluded by the quality of currently available sequencing data. This study provides the theoretical link between dN/dS values and selective coefficients in somatic evolution, and reveals the DFE for mutations in human tissues.

## Introduction

One of the principal goals of large-scale somatic genome sequencing is to uncover genetic loci under positive selection, so-called “driver” genes, that lead to clonal expansions. Enumeration of the selective advantage of each driver mutation enables prediction of future evolutionary dynamics^1^. In evolutionary biology, the distribution of fitness effects (DFE) is a fundamental entity that describes the selective consequences of a (large) number of individual mutations of an ancestral genome^2^. In somatic evolution, particularly cancer genomes, we have an extensive knowledge of the catalogue of recurrent, and likely positively selected, somatic mutations^3^, but the fitness changes associated with each mutation remain largely unquantified.

Extensive experimental effort is ongoing to determine the fitness effects of mutations. Most prominently is lineage tracing of mutations in mouse models^4,5^, but these methods are not sufficiently high-throughput to produce the DFE for all somatic mutations. Other studies have estimated the selective coefficient of somatic mutations by measuring the frequency of such mutations over time in the same individual using longitudinal sampling^6,7^ however this method is broadly limited to somatic evolution in the blood (where it is feasible to take samples from healthy individuals over time) and in rare cases of patients under active surveillance.

An alternative approach is to infer selective coefficients directly from somatic genome sequencing data. Methods to identify positively-selected (driver) mutations rely on finding genes that have significantly more mutational ‘hits’ (typically hits are non-synonymous mutations) than would be expected by chance, after correction for factors known to influence the mutation rate across the genome^8^. Conversely, negatively selected genes are expected to show a paucity of mutations^9,10^. This idea is formalised in the calculation of the dN/dS ratio – a method originally developed in molecular species evolution – that has recently been adapted for use to study somatic evolution (both cancer and normal tissue)^3,9-15^. The intuitive idea behind dN/dS is to measure the rate of non-synonymous (dN) mutations (possibly under selection) and compare that to the rate of synonymous (dS) mutations (presumed neutral). The ratio of these two numbers, each normalised for the local sequence-specific biases in the mutation rate, putatively identifies a signature of selection: dN/dS > 1 indicating positive selection, dN/dS = 1 indicating neutral evolution and dN/dS < 1 indicating negative selection.

Transforming dN/dS values to selective coefficients in somatic evolution is an unaddressed problem. dN/dS was originally developed in the context of species evolution using the Wright-Fisher process, a classical population genetics model that assumes that evolution occurs over very long timescales, which permits new mutations to fix within lineages, and also that the population size is constant, with all individuals having equal potency and non-overlapping generations. Under the Wright-Fisher model, the dN/dS of a locus is related to its selective coefficient by the relation^16^:

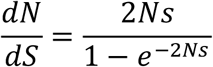

Where *N* is the effective population size and *s* the selection coefficient.

However, in somatic evolution the assumptions of the Fisher-Wright model are violated. Somatic evolution is rapid and new mutations are infrequently fixed in the population^17^, clonal dynamics are complex and population sizes unlikely to be constant^18^. Further, the lack of recombination in somatic evolution can result in strong hitchhiking effects. In addition, since in somatic evolution the ancestral genome is known it circumvents the need to measure dN/dS across a phylogeny (a necessary step for dN/dS analysis in species evolution). Violations of some of these assumptions was previously recognised to make the interpretation of dN/dS problematic^19,20^, and consequently the relationship between selective coefficients and dN/dS values is uncertain.

The size distribution of clones (called the site frequency spectrum in population genetics nomenclature) also contains information on the selective coefficients of newly arising mutations. Mathematical descriptions of the dynamics of populations of cells can make predictions on the shape of the clone size distribution under different demographic and evolutionary models^21,22^, and this approach has been used to quantify the dynamics and cell fate properties of stem cells across many tissues^23-25^. We and others have also used similar approaches to infer the evolutionary dynamics of tumours in deep sequencing data^26-29^.

To date, dN/dS analysis and the analysis of the clone size distribution have been performed independently, with conflictual results^30,31^. Here we develop the mathematical population genetics theory necessary to combine these approaches and explore how the inter-individual measure of selection at a locus as provided by dN/dS values is related to the underlying cell population dynamics that generate intra-individual clone size distributions. This approach naturally accounts for the nuances in somatic evolution that can make the interpretation of dN/dS difficult. We show how this unified approach allows for greater insight into patterns of selection than either method in isolation, and importantly reveal the precise mathematical relationship between dN/dS values and selective coefficients in somatic evolution. We use this approach to infer the selective advantage of mutations in normal tissue and examine the evolutionary dynamics of cancer subclones.

## Results

### A general approach to integrate dN/dS and clone size distributions

We present a general mathematical framework for the interpretation of frequency-dependent dN/dS values in somatic evolution. First, we construct null models of the evolutionary dynamics in the absence of selection, and then augment these models to incorporate the consequences of selection. Evolutionary dynamics differ between normal tissues and cancer cells: in normal tissues maintained by stem cells, the long-term population dynamics is controlled by an approximately fixed-size set of equipotent stem cells undergoing a process of neutral competition^32^, whereas in tumour growth the overall population increases over time. In each scenario, we develop a null model to predict the expected genetic diversity in the population in the absence of selection. Positive selection causes selected variants to rise to higher frequency than expected under neutral evolution (Figure 1a), and negative selection has the opposite effect. This insight guides how we model the effects of selection (i.e diversity of non-synonymous mutations).

**Figure 1.**
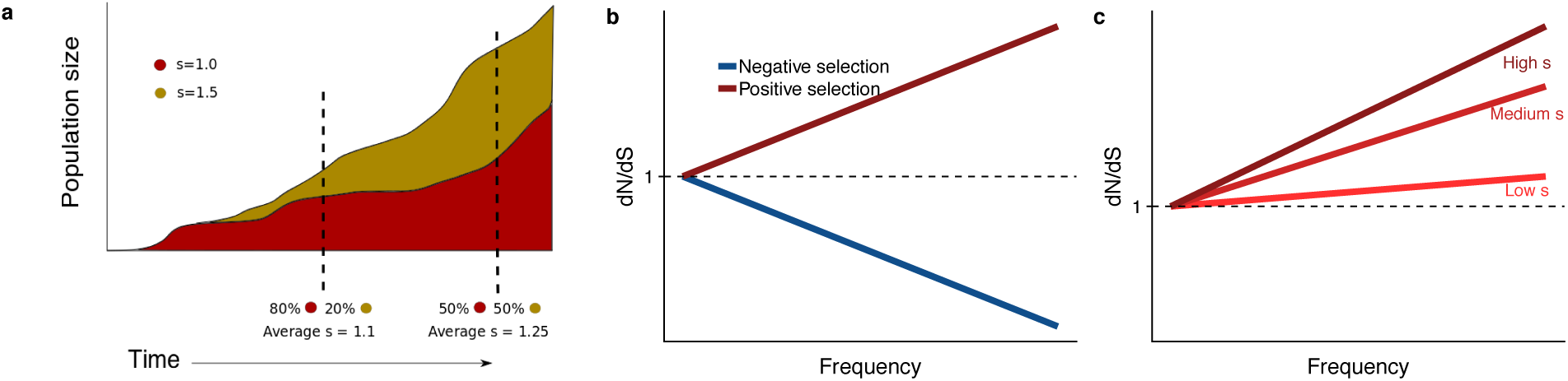
**A** Variants under positive selection are enriched at high frequency, this means dN/dS estimates are dependent on the frequency of mutation, **b.** The strength of selection influences the degree to which positively selected variants are enriched at high frequencies **c**.

Specifically, we defined the function *g*(*θ, µ, s, f*) as the expected distribution of mutations with selective (dis)advantage *s* found at a frequency *f*, for a given evolutionary dynamics scenario, where mutations accumulate at a rate *µ*. For the remainder of the paper we use passenger mutations to refer to those mutations that have no functional effect (s=0) and driver mutations those that have s>0. When comparing to data, driver mutations are taken as equivalent to non-synonymous mutations and passengers equivalent to synonymous mutations.

The functional form of *g*(*θ, µ, s, f*) encapsulates the population dynamics of the system with parameter vector *θ*, which may, for example, include the growth rate of a tumour, or loss replacement rate of stem cells in normal tissue. The direct interpretation of *s* depends on the system under question. Following the logic of the effect of selection above, for *s*′ > *s* we have that:

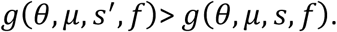

Since dN/dS measures the excess or deficiency of mutations due to selection, taking the ratio of *g*(*θ, s, m*) when *s* ≠ 0 to *s* = 0 and normalizing for the mutation rates, which may differ for passenger (*µ*_*p*_) and driver (*µ*_*d*_) mutations respectively, informs how dN/dS is expected to change as a function of the frequency *f* of mutations in the population (equation 1).

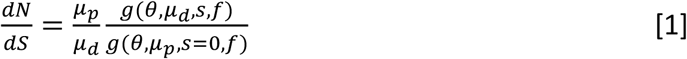

We discuss the general properties of this model. Firstly, when *s* = 0 (neutral evolution), the numerator and denominator are equal resulting in 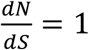, as expected. Secondly, dN/dS increases as a function of frequency *f* (clone size) for positive selection, and decreases as a function of *f* for negative selection (Figure 1b), for all *g*(*θ, µ, s, f*) that we consider. Thirdly, the shape of the curves predicted by the underlying population model encodes the value of the selection coefficient; for example the steepness of the increase is proportional to the selection coefficient *s* (Figure 1C). These observations are a natural consequence of positive selection driving selected mutations to higher frequency (Figure 1a).

Unfortunately, directly using equation [1] to measure selective coefficients from the slope of the dN/dS curve as function of frequency is often impractical. Real sequencing data often suffers from a limited number of mutations detected at any particular frequency and measurement uncertainties in these frequencies. To circumvent these issues, we introduce “interval dN/dS” (i-dN/dS) that aggregates over a frequency range to reduce the influence of these sources of noise. Interval dN/dS is defined as:

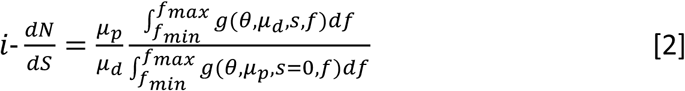

Fixing the integration range [*f*_*min*_, *f*_*max*_] allows for robust inference of *s* in potentially sparse and noisy sequencing data using maximum likelihood methods (see Methods).

### Frequency-dependent dN/dS values in stem cell populations

In healthy tissue, only mutations that are acquired in the stem cells will persist over long times, and so we restrict our attention to these cells. Quantitative analysis of lineage tracing data has shown that the stem cell dynamics of many tissues conform to a process of population asymmetry^32^. In this paradigm, under homeostasis, the loss of stem cells through differentiation is compensated by the replication of a neighbouring stem cell, thus maintaining an approximately constant number of stem cells. These dynamics are represented by the rate equations:

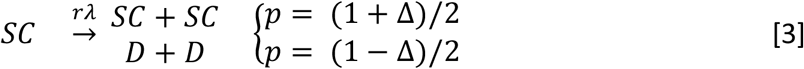

where *SC* refers to a single stem cell which divides symmetrically to produce either two stem cells or two differentiated cells (denoted as *D* above), *λ* is the rate of cell division per unit time, and *r* is the probability of a symmetric divisions. The product *rλ* is referred to as the loss/replacement rate. Differentiated cells will ultimately be lost from the population over long time scales. Under homeostasis, these processes should be exactly balanced with Δ= 0. With Δ≠ 0, the fate of a stem cell is ‘biased’, introducing positive or negative selection into the model. Previous mathematical analysis shows that this model is a good description of the clonal dynamics in the oesophagus and skin^23,33,34^. Using the previous analytical results describing the temporal evolution of the clone distribution (see supplementary methods for detailed discussion) we derive the frequency distribution *g*(*θ, µ, s, f*) for oesophagus and skin as ^21,23,35^:

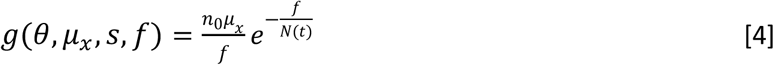

Where *n*_0_ is the starting population size and *µ*_*x*_ the mutation rate, which may be different for drivers (*s* ≠ 0) and passenger mutations (*s* = 0). *N*(*t*) is a scaling factor that depends on Δ, the bias toward self-renewal, which we interpret as our selection coefficient in this system. Specifically:

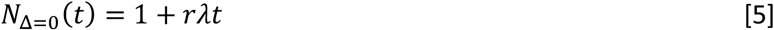

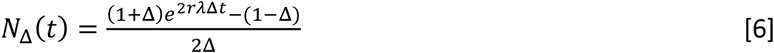

We note that at long times (large *N*(*t*)) equation [4] converges to a 1/*f* distribution for the site frequency spectrum of a fixed size population^36^. *N*(*t*) can be interpreted as the average size of a labelled clone after time *t*, which even under homeostasis grows over time and compensates for some clones being lost due to drift. From these expressions, we can then write down a closed-form expression for i-dN/dS as a function of clone frequency (see methods) that allows for maximum likelihood estimation of parameter values (Δ, *rλ*). We confirmed the accuracy of our derivation using simulations (Figure 2a), and performed power calculations to determine the minimum number of mutations required to correctly infer the underlying population dynamics. We determined that 8 mutations per gene was sufficient to accurately recover Δ (Figure 2b) with accuracy increasing for higher mutation burdens (Figure 2c).

**Figure 2.**
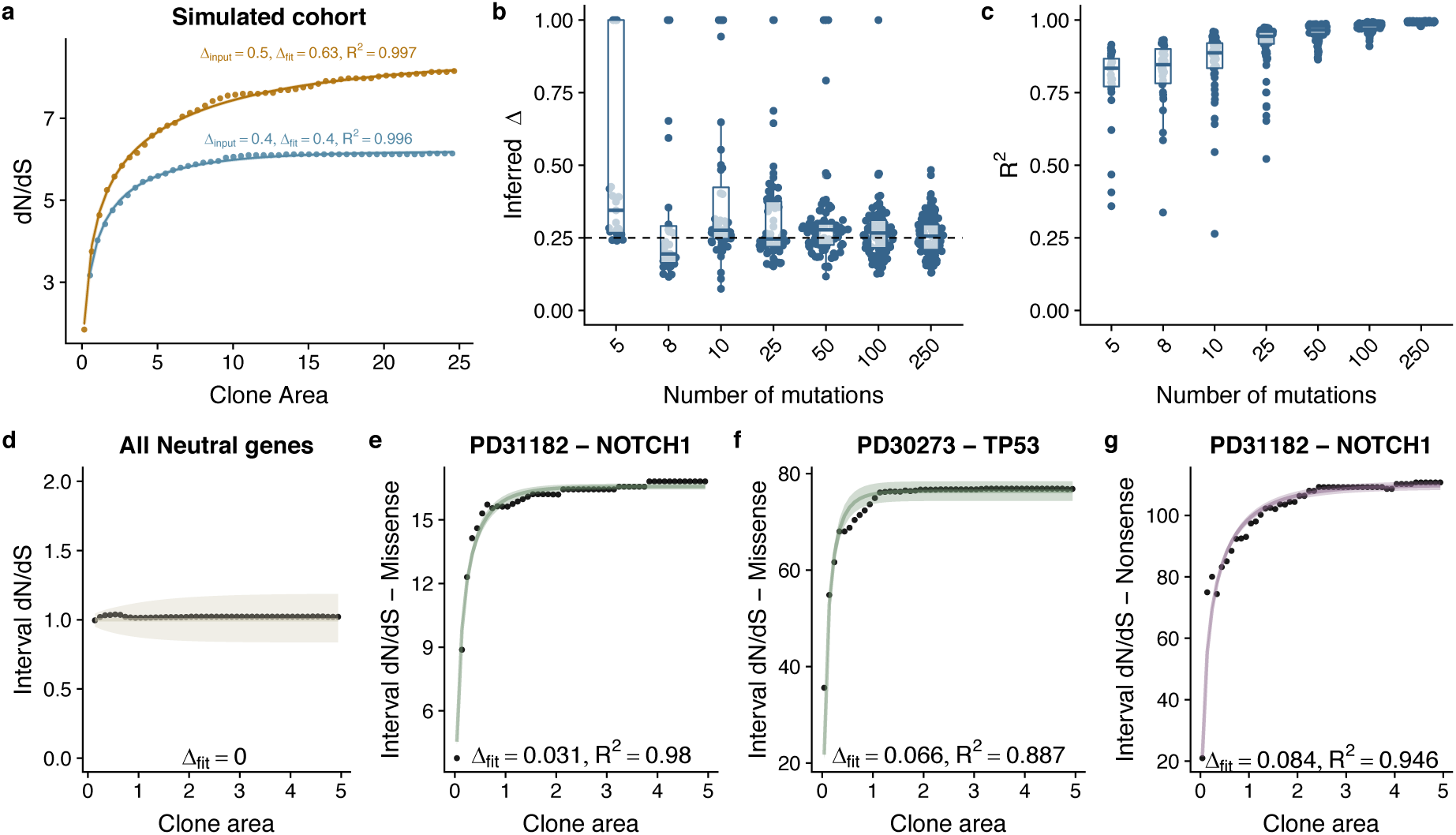
**a** Interval dN/dS as a function of clone area for 2 simulated cohorts where driver mutations induce different biases, theoretical model captures the dynamics well and enables us to recover the bias Δ, accurately. As the number of mutations increases ability to recover the correct Δ and the model fit (measured using R^2^) improves **b** and **c. d** Data and model fit for all neutral genes, shows i-dN/dS = 1 across the frequency range and inferred bias of 0. Data and model fit for **e** NOTCH1 missense mutations in patient PD31182, **f** missense TP53 mutations in PD30273 and NOTCH1 nonsense mutations in PD31182. Data are black points and model fits are solid lines with shaded areas denoting 95% CI.

### Selection advantages in histopathologically normal human oesophagus

We inferred the selective advantage of driver mutations in human oesophagus using published deep sequencing data from Martincorena and colleagues^14,37^ that documents the clonal expansion of a panel of putative driver mutations in histopathologically-normal oesophageal biopsies.

We used the dndscv bioinformatics tool^3^ to calculate frequency-dependent dN/dS values from these data (clone size measured in fraction of mutant reads multiplied by 2mm^2^ – the area of the biopsy – and assuming 5,000 stem cells per mm^2^ tissue). dN/dS values varied considerably as a function of mutation frequency (Figure S1).

We considered the average frequency-dependent dN/dS values across all genes in the panel, on a patient-by-patient basis. Our theoretical model of i-dN/dS calculated from these data fitted strikingly well (Figure S2). Estimates of the loss/replacement rate *rλ* of the stem cell population were in the range 1.2-5.0 per year (Figure S2&S3). Inference of the selective advantage *s* (measured in terms of the bias towards self renewal Δ) revealed an average bias of 0.004 (0.002 – 0.005 95% CI) per missense mutation (Figure S2). Nonsense mutations caused a five-fold greater bias towards self-renewal of 0.021 (0.008 – 0.032 95% CI) (Figure S3). After removal of all genes that are strongly selected, global dN/dS values on the remaining 48 genes show dN/dS of approximately 1 across the frequency range (Figure 2d), and i-dN/dS analysis revealed somatic mutation does not associate with a proliferative bias (Δ=0).

We then fitted the data on a gene-by-gene and patient-by-patient basis for cases where sufficient mutations were available to perform the fit (Figure 2e-g; Figure S4). A broad range of selective advantages were inferred (Figure S4&S5). Mutations in *TP53* showed large biases across all patients for both missense, Δ=0.057 (0.05-0.068 95% CI) and nonsense mutations, Δ=0.094 (0.091-0.097 95% CI) (Figure 3a-b). This was also true for mutations in NOTCH1 with Δ=0.029 (0.019-0.036 95% CI) for missense and Δ=0.072 (0.034-0.089 95% CI) for nonsense mutations. *NOTCH2, PIK3CA, CREBBP* and *FAT1* also showed a bias toward self-proliferation in multiple patients (Figures 3a-b), though most had a small effect on fitness (range 0.003 – 0.029 for missense mutations and 0.030 – 0.041 for nonsense mutations). Together these data suggest a distribution of fitness effects (DFE) characterized by many small effect mutations with few large effect mutations (Figures 3c-d), as in seen in organismal evolution^2^.

**Figure 3.**
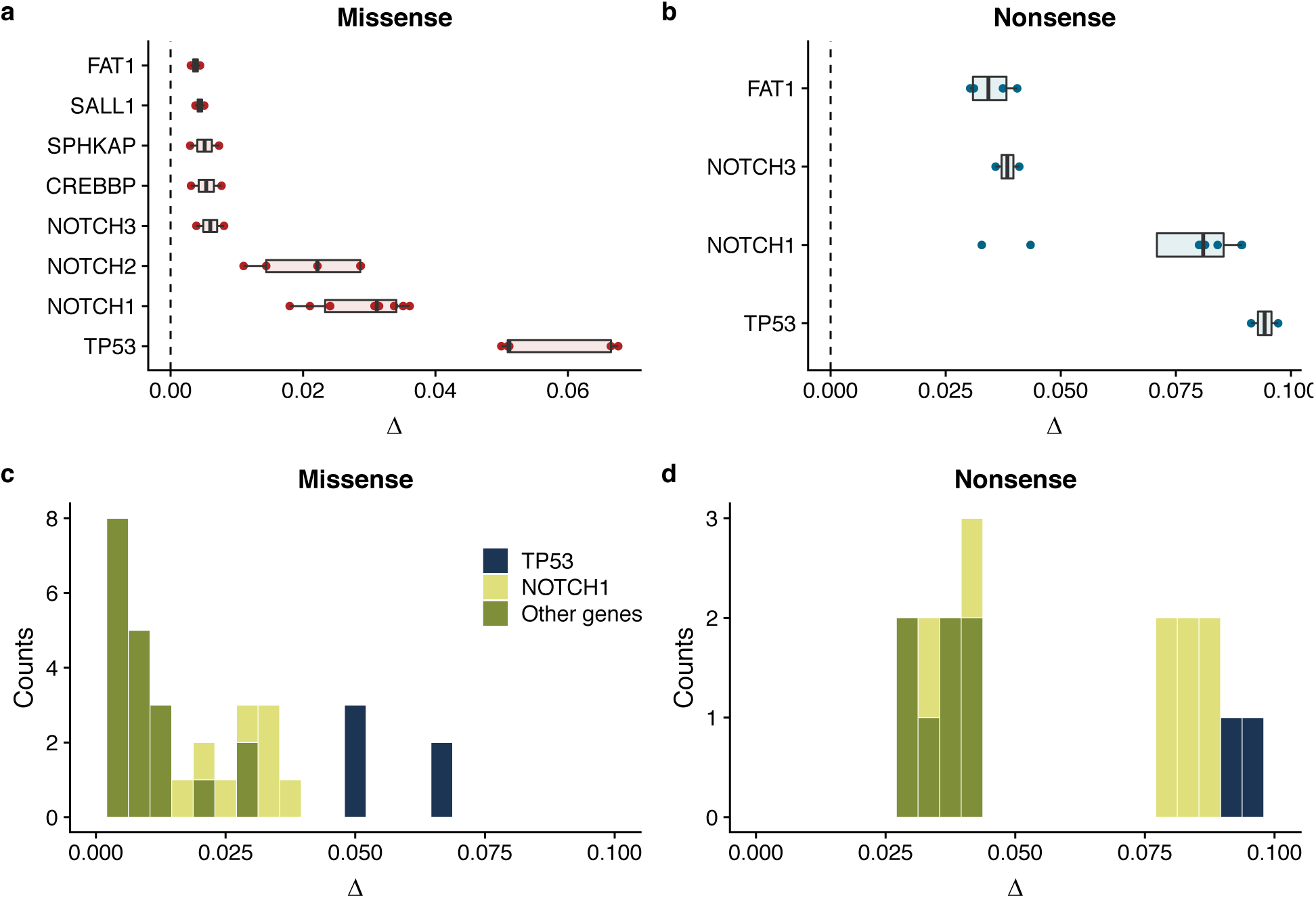
Summary of model fits across all patients for normal oesophagus data. Inferred biases Δ for genes where at least 2 patients had good model fits (R2 > 0.6 & >7 mutations) for missense mutations **a**, and nonsense mutations **b**. Inferred distribution of fitness effects for all genes across all patients for missense mutations **c**, and nonsense mutations **d.**

### Driver mutation selective advantage in normal skin

Martincorena and colleagues have also published data on the expansion of driver mutations in ostensibly normal human skin^18^. Analyses of these data with interval dN/dS revealed a per-patient average selective advantage per mutation (again measured in terms of the bias towards self renewal Δ) of Δ=0.001 for missense mutations and four-fold higher for Δ=0.004 for nonsense mutations (Figures S6a-c). Performing the analysis on a gene-by-gene basis was limited by the low detected number of mutations, and the limited frequency range (clone size range). Good fits to the data were obtainable for *NOTCH1* missense mutations in patient PD18003 with fitness estimated to be Δ=0.0149 (0.0148-0.0150 95% CI), and TP53 missense mutations also in patient PD18003, Δ=0.0054 (0.0051-0.0058 95% CI) Figure S6. These fitness coefficients were similar to the oesophagus data. For missense mutations we were also able to produce the distribution of fitness effects across the skin cohort, which showed similar characteristics to the oesophagus data of a small number of high effect mutations and a larger number of smaller effect mutations, Figure S6f.

### Clonal mutations have greater dN/dS than subclonal mutations in cancers

We next investigated the selective advantage of driver mutations in cancer. We first investigated whether or not differences existed between dN/dS values for clonal mutations (ie truncal, present in all cells in a cancer) and subclonal mutations (present in a subset of cells in a cancer) were apparent. Using sequencing data from 2,619 cancers from TCGA that had sufficient cellularity and depth (see Methods) we calculated the mutation copy number (MCN) for each mutation and grouped mutations into subclonal, clonal and amplified across the cohort, where mutations with MCN < 1 were subclonal, MCN == 1 were clonal and MCN > 1 were amplified (Figure 4a). We than calculated global dN/dS ratios for a panel of 198 high confidence driver genes (Methods).

**Figure 4.**
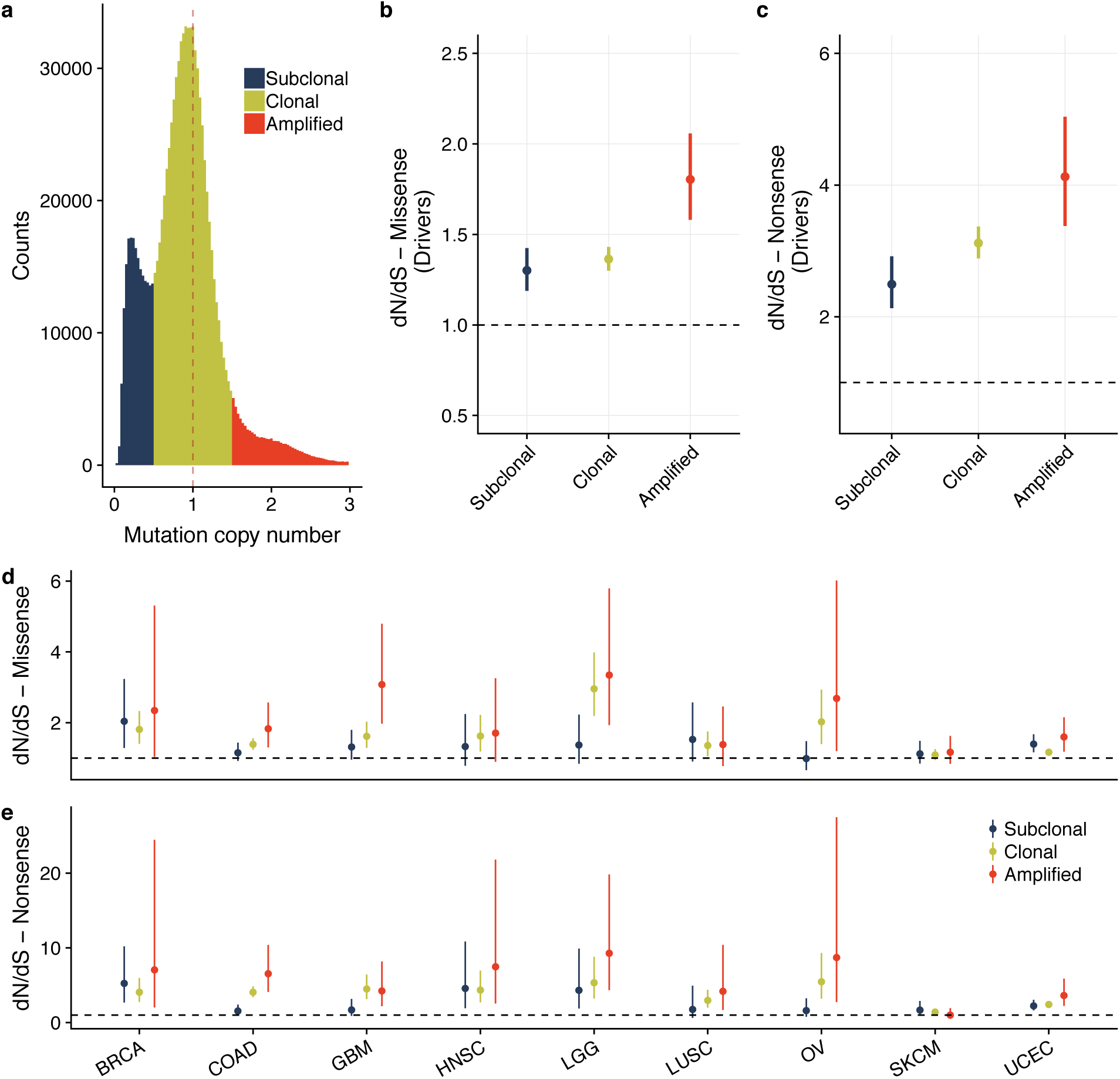
Mutation copy number histogram across 2,619 TCGA samples coloured by mutation clonality, **a**. dN/dS by mutation clonality for missense, **b** and nonsense **c** mutations in a panel of 192 high confidence driver genes. The same analysis done per cancer type for missense **d** and nonsense **e**.

Across all cancers, the signal of positive selection was more pronounced for clonal mutations (Figures 4b-e), with the highest dN/dS values found in amplified mutations^38^. Subclonal mutations on the other hand demonstrated much lower dN/dS values. The same pattern was also evident in individual cancer types (Figure 4e,d & S7). In many cancer types (colorectal, ovarian, glioblastoma) subclonal mutations showed no evidence of subclonal selection (neutral evolution; dN/dS = 1), Figure 4e,d & Figure S7.

### Interval dN/dS for cancer

We applied our mathematical approach above to calculate i-dN/dS in cancer evolution. In cancer evolution *g*(*θ, µ, s, f*) must account for tumour growth dynamics and subclonal mutations which may rise and fall in frequency due to selection and drift. The well-studied Luria-Delbrück distribution and its extensions describes these dynamics^39^. Specifically, the Luria-Delbrück distribution describes the expected number of mutational lineages at a particular frequency assuming an underlying birth-death process for individuals in the population. For neutral mutations the site frequency spectrum has a characteristic 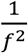 dependence, where *f* is the frequency of the mutations ^35,40^. Hence:

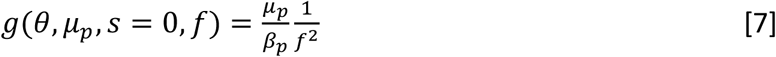

where *µ*_*p*_ is the passenger mutation rate and *β*_*p*_ is the survival probability of a lineage at division. We previously showed that in many cancers across types (approx. 30% of cases), subclonal mutations closely follow the prediction of this neutral model^26^.

Extensions to the classic Luria-Delbruck distribution describe the differential fitness of mutants. We defined the relative fitness advantage *s* as the ratio of net growth rates between wildtype ‘passenger’ mutations (*λ*_*p*_) and driver mutations (*λ*_*d*_) :

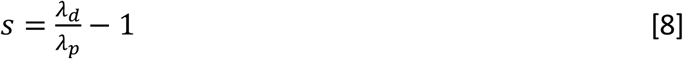

s > 0 indicated positive selection while s < 0 indicated negative selection. We also defined the birth and death rates of the respective wildtype (passengers) and mutants (drivers) as *b*_*p*_, *d*_*p*_, *b*_*d*_ and *d*_*d*_. Here, the site-frequency distribution again follows a power law but with exponent dependent on the relative fitness advantage of the mutant ^35,40^:

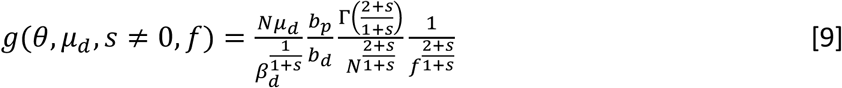

Here, N is the tumour population size at the time of sampling. Using these expressions (equations 7&9), we derive i-dN/dS (see Methods). The equation exhibits the same qualitative behaviour as for the stem cell model, in that dN/dS increases as a function of frequency for positive selection and decreases for negative selection (Figure 5a). Using a simulation-based model to generate synthetic data, we confirmed the accuracy of the model by accurately recovering the inputted selection coefficient by application of the theoretical model and maximum likelihood inference (Figure 5a).

**Figure 5.**
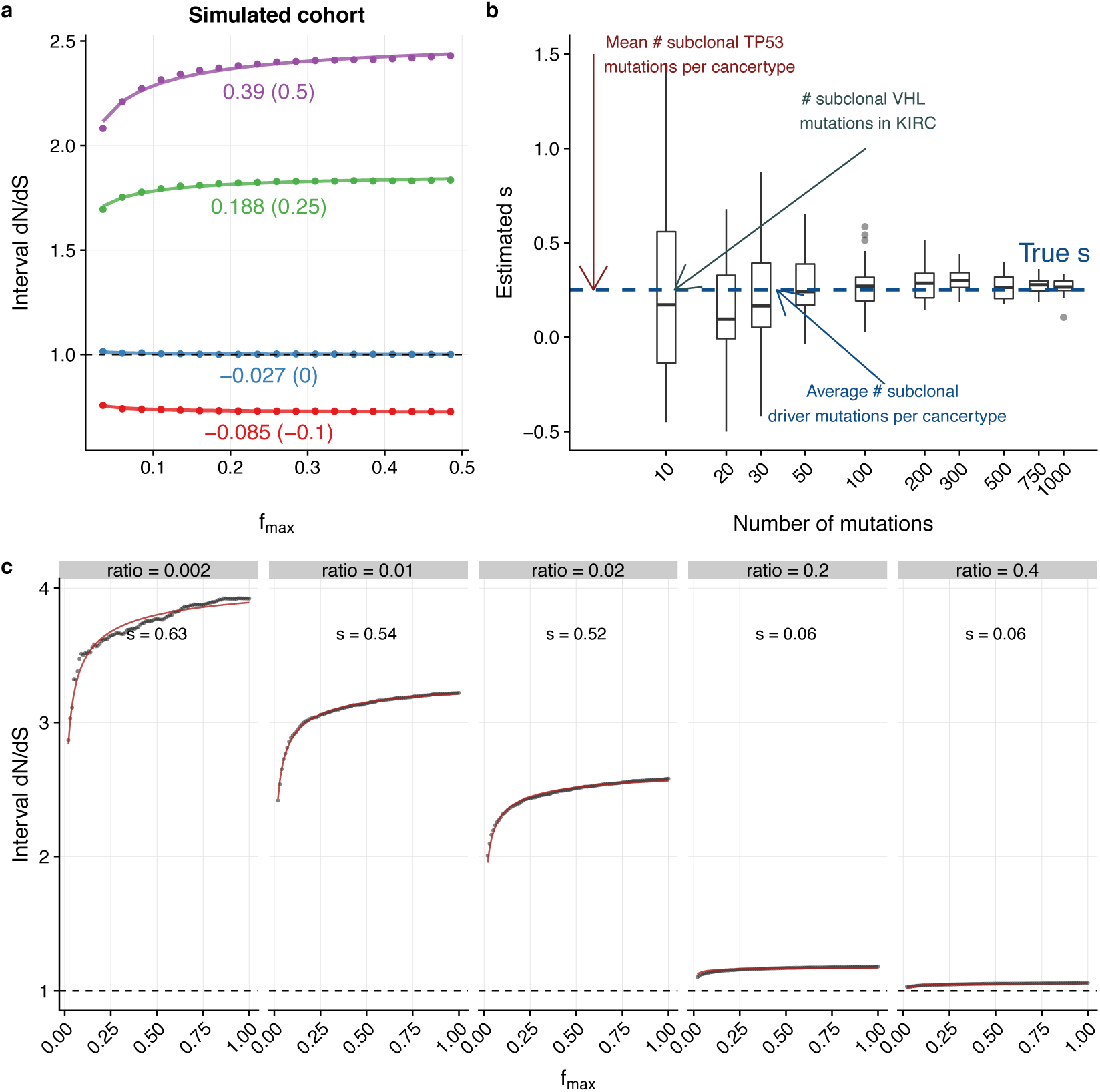
Interval dN/dS as a function of frequency for 4 simulated cohorts where driver mutations induce different selective advantages, **a**. Points are simulated data and lines are model fits, under each line is the inferred selective advantage and the true selective advantage in brackets. Power to correctly infer the selection coefficient depends on the number of mutations in the cohort, **b**. We generated a cohort of 1000 tumours and then subsamplesd the mutations (50 times) and inferred the selection coefficient. For TCGA we are limited by a small number of subclonal drivers to accurately perform the inference. The ratio of the driver mutation rate to passenger mutation rate has a strong influence on dN/dS, **c**. Here we generated synthetic cohorts where the strength of selection of driver mutations was 0.5, and different ratio of driver mutation rate to passenger mutation rate. When drivers are rare, dN/dS > 1 and we can accurately apply our model. When drivers are frequent compared to passengers we observe strong hitchhiking effects which results in dN/dS∼1.

Subclonal dN/dS is strongly influenced by the ability to resolve low frequency variants. We generated synthetic tumour cohorts that modelled subclonal selection, and simulated ‘perfect sensitivity’ for mutation detection. In these cases, where all mutations were resolved, we measured dN/dS≈1 (and hence infer a selection coefficient of 0), despite some lineages being positively selected (Figure S9). If only higher frequency variants were analysed, then the measured dN/dS > 1 and the correct selective coefficient is inferred (Figure S9). We note that at very low frequencies the detected mutations are newly arisen in the population, and so are as yet ‘unfiltered’ by selection. Consequently the ratio of non-synonymous to synonymous mutations is expected to be proportional to the respective mutation rates of the two mutation types. The abundance of low frequency mutations also increases exponentially with decreasing clone frequency, and so including very low-frequency variants ‘drowns out’ the effects of selection (Figure S9C). We note that the limited sequencing data of the majority of currently available cancer genomic data means that typically only high frequency variants are detected.

### Currently available cancer sequencing data is insufficient to infer selective advantages

Limitations in the quality of currently available sequencing data meant that the theoretically predicted frequency dependence of dN/dS values could not be assessed in cancer genomics data (Figure S8). Limited sequencing depth introduces uncertainty into the determination of variant allele frequencies (“sequencing noise”) which can result in incorrect classification of mutation clonality. Visual inspection of the mutation copy number histogram for TCGA data (Figure 4a) showed a very broad dispersion of MCNs, and the resolution at lower (subclonal) frequencies was particularly poor. Issues arising due to sequencing noise are exacerbated in the setting of dN/dS analysis where pooling the data from multiple patients with different sequencing depth and purities is required. Consequently, the range of subclonal frequencies where interval dN/dS could be calculated was severely restricted.

We tested whether or not looking at individual genes (rather than individual mutations) allowed for measurement of the DFE. However, the lack of recurrent subclonal mutations on a gene-by-gene basis precluded this approach. Power calculations predicted that a minimum of 30 subclonal mutations in a given gene were required to accurately fit the interval dN/dS model (Figure 5b). This level of subclonal recurrence of individual mutations was not seen in the data: for example, the average number of subclonal mutations in *TP53* per cancer type, as well as the number of subclonal VHL mutations (which has been reported to occur subclonally at an appreciable frequency ^41^) were both well below this cutoff (Figure 5B). Consequently, large cohorts of tumours sequenced to higher depth are required to apply this approach.

Aside, we note that the traditional dN/dS approach, and also our modelling framework, assumes that mutations are independent, and consequently the possibility of hitchhiking of mutations (e.g. nested driver mutations within clones) is neglected. In simulated data, we observed high mutation rates for both driver and passenger mutations led to hitchhiking being common, and subsequent obscuring of the signal of selection (Figure 5c). In extreme cases this led to dN/dS = 1 (apparent neutral evolution) even in the presence of multiple selected lineages. For most cancers, the number of driver mutations per cancer is thought to be low (<10)^3^, but nevertheless in hypermutator cancers the hitchhiking effect is likely to be common. Thus, despite hypermutator tumours tending to have fewer copy-number alterations and hence less problematic estimation of MCNs, the prevalence of hitchhiking precludes analysis of these tumours.

## Discussion

Here we have shown that the combination of dN/dS values with mutation frequency-based information provides additional quantitative insight into dynamics of somatic evolution than either method alone. Specifically, the combined approach enables direct inference of the selection coefficients of mutations in somatic tissues.

Using this methodology we have begun the construction of the distribution of fitness effects (DFE) in somatic evolution (Figure 3c,d & Figure S6f). In histologically normal epithelium, mutations of most genes considered showed minimal effects on fitness (near-neutral evolution), though selection coefficients for some loci, foremost *NOTCH1* and *TP53* were considerable (>1% and >5% respectively), and consequently the DFE has most mass close to s=0 with a long right-tail of highly-selected variants. We observed that values of selective coefficients of individual genes varies between patients, likely because of inter-patient difference in the precise location of point mutations, but potentially also because of inter-patient variation in selective pressure from the microenvironment. Nevertheless, the comparative rank of per-gene fitness coefficients was broadly consistent across patients. This consistency in selective coefficients is in agreement with the observation highly recurrent gene mutations in cancer^42^ and evidence of repeatability in cancer evolution^43^.

We have previously measured fitness effects in individual cancers (but were unable to ascribe fitness changes to individual genes) finding increases in growth rate in a selected clone approaching 100% in some cases^27^. Care must be taken when comparing selective coefficients between normal and cancer populations, because in the former we quantify selection as tilt away from homeostasis and towards net growth of a lineage, whereas in cancer we infer the relative growth rate of a clone within the tumour as a whole. With this important caveat in mind, nevertheless the fitness increases observed in cancer appear to be much larger than for normal tissues. We hypothesise that this is because the effect of selection is weaker in expanding populations like cancer, wherein the generation of a subclonal expansion requires very large increases in fitness^44^.

On a cautionary note, our theoretical work shows that the clonality of mutations strongly determine the observed value of dN/dS, and so a misleading picture of the selective forces operating in a tumour (or healthy tissue) will be produced if dN/dS frequency-dependent effects are not corrected for. The accuracy of any estimate of evolutionary dynamics from dN/dS values is of course dependent of the underlying accuracy of the dN/dS measure itself, which is compromised by uncharacterised variability in the mutation rate across the genome^45^ and in the uncertain pathogenicity of individual single nucleotide variants (extensions to estimate site level selection coefficients may circumvent some of these issues^46,47^). Finally, we note that dN/dS measures cannot elucidate evolutionary pressures in individual samples as insufficient (subclonal) mutations will be found at any individual locus. dN/dS cohort measurements are sensitive to outliers, where a few patients with high selection can drive the results ^48^. Other approaches, such as using the site frequency spectrum, are likely more powerful for these types of questions.

Combining population genetics methods with comparative genomics is a powerful way to infer selection pressures in human somatic evolution, giving new insight into the fundamental parameters that determine evolutionary dynamics in health and disease.

## Acknowledgements

A.S. is supported by the Wellcome Trust (202778/B/16/Z) and Cancer Research UK (A22909). T.G. is supported by the Wellcome Trust (202778/Z/16/Z) and Cancer Research UK (A19771). We acknowledge funding from the National Institute of Health (NCI U54 CA217376) to A.S and T.A.G. This work was also supported a Wellcome Trust award to the Centre for Evolution and Cancer (105104/Z/14/Z). C.P.B. acknowledges funding from the Wellcome Trust (209409/Z/17/Z).

## Methods

### TCGA Data Processing

MAF (Mutation Annotation Format) files from the Mutect2 mutation calling algorithm and copy number segmentation data for 9950 cancers from 26 cancer types were downloaded from the genomic data commons portal using the TCGAbiolinks R package ^49^. Cellularity and ploidy estimates derived from ASCAT were obtained from COSMIC (https://cancer.sanger.ac.uk/cosmic/download). We then filtered for >2 reads reporting the variant and >9 reads coverage at each locus in both the tumour and normal sample. We removed samples where the effective depth (defined as cellularity times depth) was < 50X and those that had likely undergone genome doubling (ploidy > 2.5). This left 2619 samples from 17 cancer types which we deemed suitable for analysis.

Copy number (CN) segmentations together with cellularity estimates were used to correct the variant allele frequency and produce mutation copy number estimates. We assume that the observed CN state 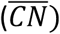 was a combination of signals from the tumour sample and contamination from normal cells (with two copies) assuming tumour purity c.

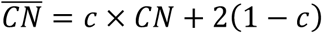

With this, log(R) ratios were transformed into copy number states using the following formula:

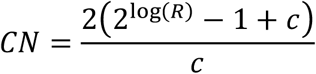

Using these corrected copy number states, mutation copy number (MCN) values were calculated. Given mutation *i* with variant allele frequency *VAF*_*i*_, copy number *CN*_*i*_ at the locus and cellularity estimate of the tumour c, the MCN was calculated as follows:

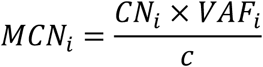

Visual inspection of the MCN histograms (Figure 4a) show a dominant peak at MCN = 1 representing clonal mutations present in a single copy, confirming that the corrections we applied work as intended.

### Oesophagus and skin data

For the oesophagus and skin data we used mutation calls provided by the original studies. In the oesophagus data when a mutation was present in multiple adjacent biopsies we used the sum of the mutation frequency times the area of the biopsies (2mm^2^) as our readout of clone size and performed the dN/dS analysis on a patient by patient basis.

### dN/dS calculations

For calculating dN/dS ratios the dndscv R package was used which calculates both global dN/dS ratios across the whole exome or a panel of genes as well as per gene dN/dS ratios using a covariate based model to infer dN/dS values with a limited number of mutations ^3^. In an attempt to enrich for positive selection in some of our analysis we calculated dN/dS for a subset of 198 high confidence driver genes ^50^.

Over or under filtering of possible germline SNPs is known to influence dN/dS values in somatic genomes^3^. We previously found that mutation calls provided by TCGA are likely over stringent on filtering germline SNPs resulting in inflated dN/dS values ^48^. To circumvent this issue, we calculated a baseline dN/dS value by randomly selecting 1,000 genes (excluding drivers) and then running dndscv across the whole TCGA cohort, reasoning that this should on average return dN/dS = 1, and any deviation from this would be due to under/over filtering of SNPs. Repeating this procedure 50 times and then taking the mean value gave us our baseline value which we could then subtract from further dN/dS values we calculate in our analysis. To confirm this procedure produces the expected result of dN/dS = 1 in the absence of selection, we repeated the procedure and again, randomly selected 1,000 genes 100 times and then applied the correction (subtracting the calculated deviation from 1). As would be expected the mean of this distribution was dN/dS = 1, validating our approach, Figure S10.

To calculate the interval dN/dS measure we took our corrected mutation frequency data and determined a low cutoff *f*_*min*_ based on the minimum mutation frequency. We then created a vector of frequencies *f*_*max*_ that covered the total range of mutation frequencies and calculated dN/dS between *f*_*min*_ and all values of *f*_*max*_. This allowed us to plot dN/dS as a function of *f*_*max*_ and fit our interval dN/dS models.

### Model fitting

We used a maximum likelihood approach to fit our models to the data. Defining the observed interval dN/dS as *y* and the model dN/dS as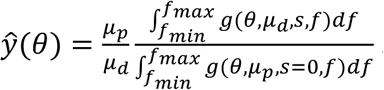. First of all we define the residuals between the data and the model as *R* = *y* − *ŷ*. Assuming that the residuals are normally distributed with mean 0 we can write down the negative log likelihood (NLL) as

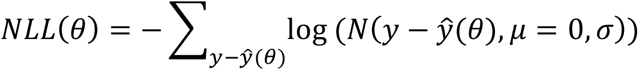

where *N* denotes the normal probability density function. We can then find the parameters *θ* that minimize the NLL and calculate confidence intervals on these estimates using the Fisher information matrix.

### Interval dN/dS models

For the stem cell model, using equations [2]-[6] in the main text, interval dN/dS is given by:

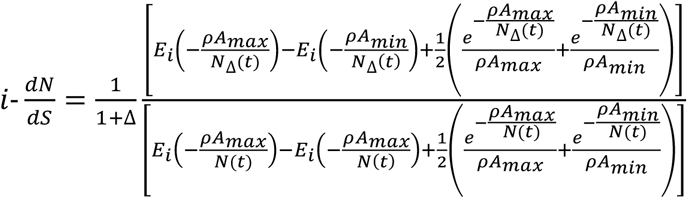

Where *E*_*i*_ is the exponential integral 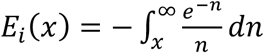. Given that the data is in terms of area, A we made the transformation *f* = *ρA*, where *ρ* is density of stem cells per mm^2^, which we set to 5,000 cells /mm^2^ for fitting.

For the cancer model, interval dN/dS is given by:

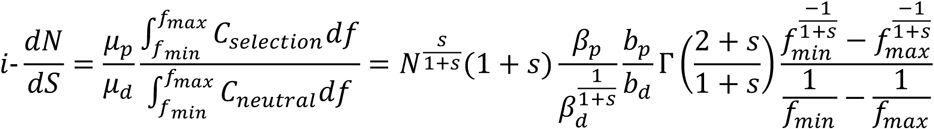

We note that in the cancer setting because the final population size N is generally unknown we fit the model 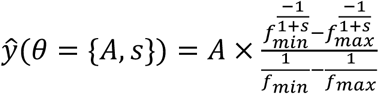

For a detailed description of the mathematical background of the clone size distribution in these models and comparison with simulation see the supplementary Jupyter notebooks.

### Simulations

To confirm our analytical models and investigate the influence of uncertainty in mutation frequencies due to sequencing noise and to challenge some of the underlying assumptions of our theoretical approach, we developed 2 simulation based models. The first one models cancer evolution and the second models stem cell evolution under homeostasis. For the cancer evolution model, we adapted our previously described model^27^ so that mutations can be one of two types, neutral passengers or mutations that have an effect on fitness of cells (either positive or negative). We model cancer growth as a continuous time branching process. At each division, daughter cells acquire mutations with a fitness effect s at rate *µ*_*d*_ and passenger mutations (which are neutral) at rate *µ*_*p*_. This is implemented by drawing a Poisson random variable with mean given by *µ*_*d*_ or *µ*_*p*_. Fitness of passenger mutations is 0, while driver mutations have fitness advantage s, where s is defined by equation [8]. We also implemented a model where fitness was a random exponentially distributed variable with mean s.

For the stem cell model we seed a population of *N*_*s*_ stem cells that then undergo loss/replacement as described by the following rate equations

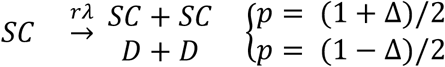

As only the stem cells are long lived the differentiated cells are not explicitly modelled such that when a stem cell “differentiates” it is effectively lost from the population. As in the cancer model, during division, daughter cells acquire mutations with a fitness effect at rate *µ*_*d*_ and passenger mutations at rate *µ*_*p*_. Fitness increases the bias toward self-proliferation Δ of a stem cell lineage. Additional driver mutations do not further increase the fitness of stem cells.

To calculate dN/dS across a cohort of simulated tumours or tissue biopsies we count the number of driver mutations *N*_*d*_ and the number of passenger mutations, *N*_*p*_ and then normalize by their respective mutation rates. In our model drivers = non-synonymous and thus every driver has an effect on fitness. Then the ratio of these two numbers gives us the excess or deficit of mutations due to selection – ie the dN/dS ratio.

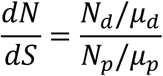

For the interval dN/dS we simply calculate the *N*_*x*_ between *f*_*min*_ and *f*_*max*_.

To introduce uncertainty into mutation frequencies we perform a process of empirically motivated sampling to the true underlying frequency *f*. Firstly, we specify the average depth of sequencing D, then the depth of sequencing for mutation i is given by

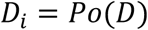

The sampled number of read counts is then

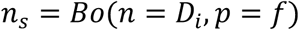

And the sampled variant frequency is then *f*_*s*_ = *n*_*s*_/*D*_*i*_

### Code and data availability

Code used for the analysis are included as a snakemake pipeline which will reproduce all the analysis and generate all the figures. Julia ^51^ was used for the majority the simulations and R ^52^ was used to analyse the data and generate the figures. Some of the analysis rely in bespoke packages written for this which are freely available under and open source licence. Code is available at github.com/marcjwilliams1/dnds-clonesize.

---

**Figure S1.**
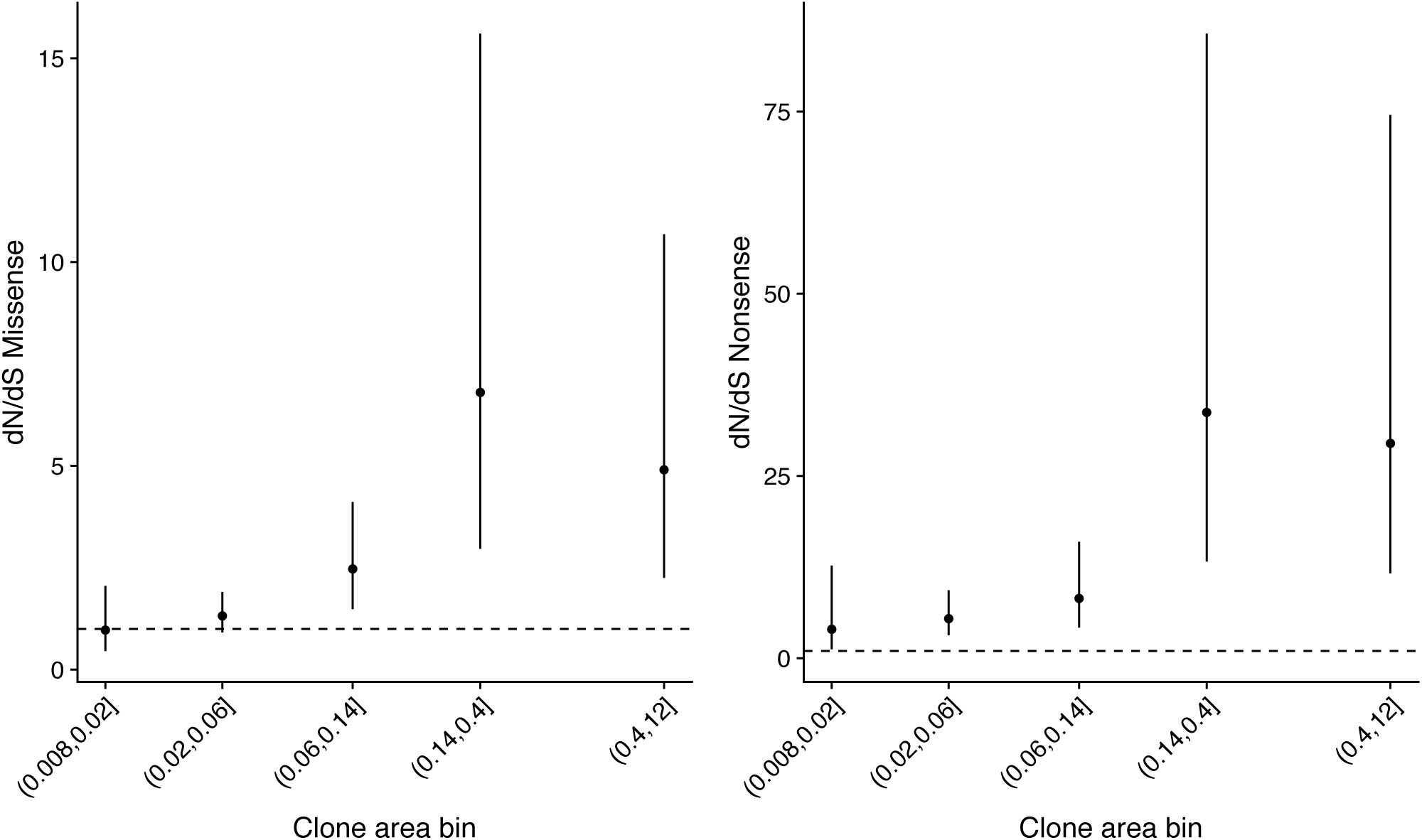
Global dN/dS values in different frequency bins for patient PD31182 showing that the values depend on the frequency of mutations.

**Figure S2.**
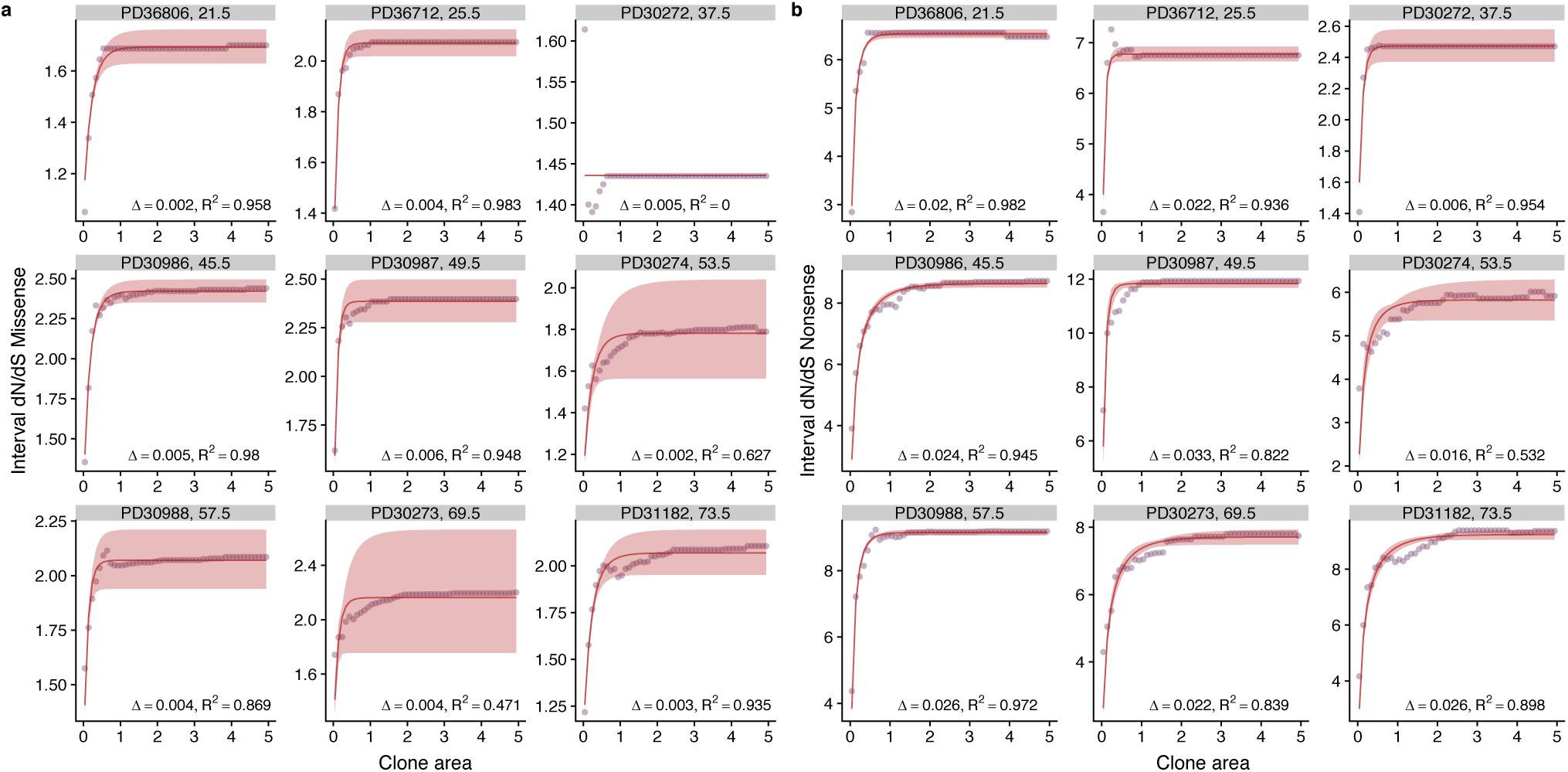
Model fits for all patients in the oesophagus data set. Purple points are data and red lines model fits. Fits were performed separately for missense, **a** and nonsense mutations, **b.** Each plot is annotated with the inferred bias Δ and the R^2^ value.

**Figure S3.**
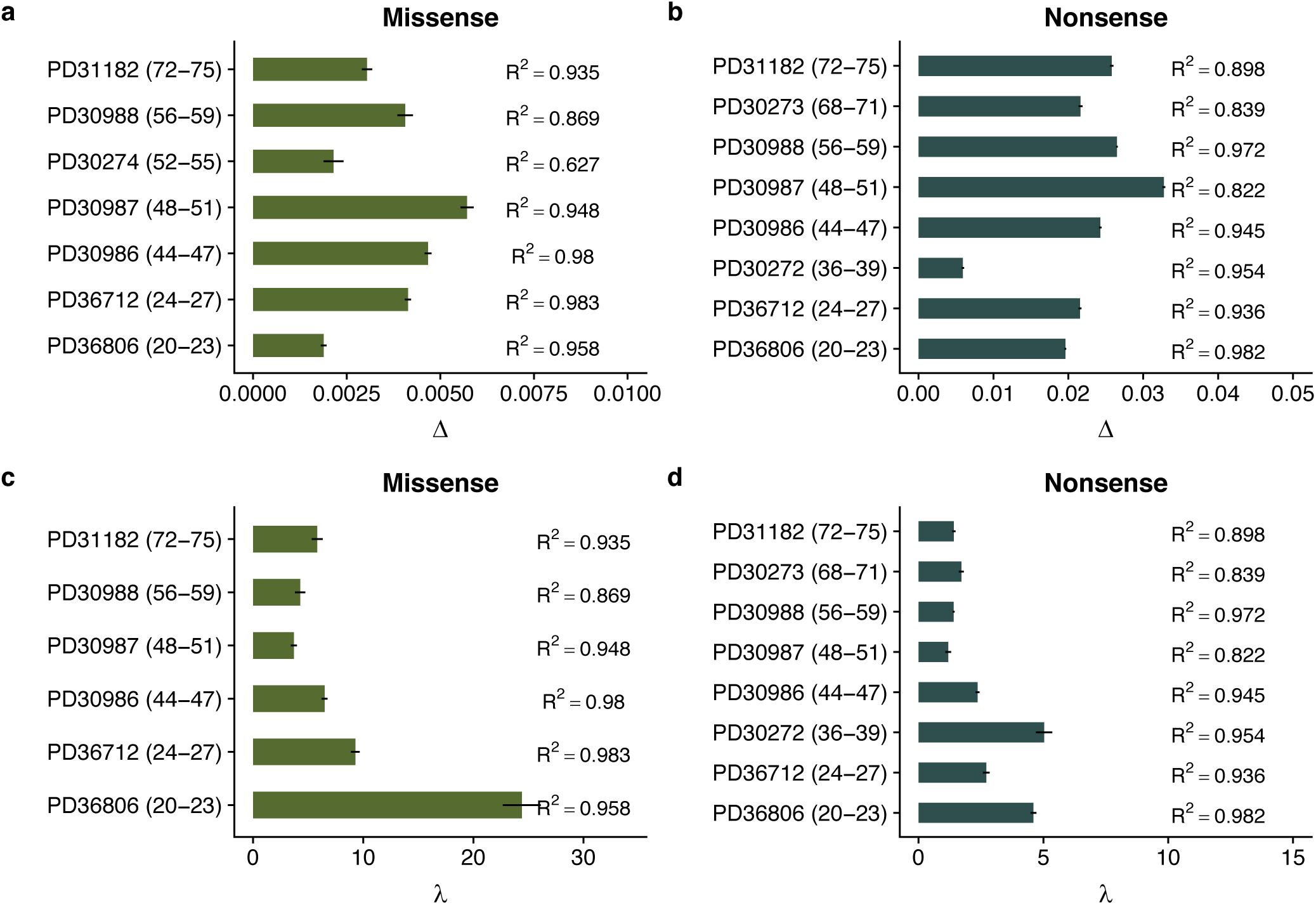
Inferred biases for for each patient in the oesophagus dataset based on missense, **a** and nonsense mutations, **b**. Inferred loss replacement rates, *λ* for each patient based on missense, **a** and nonsense mutations, **b**.

**Figure S4.**
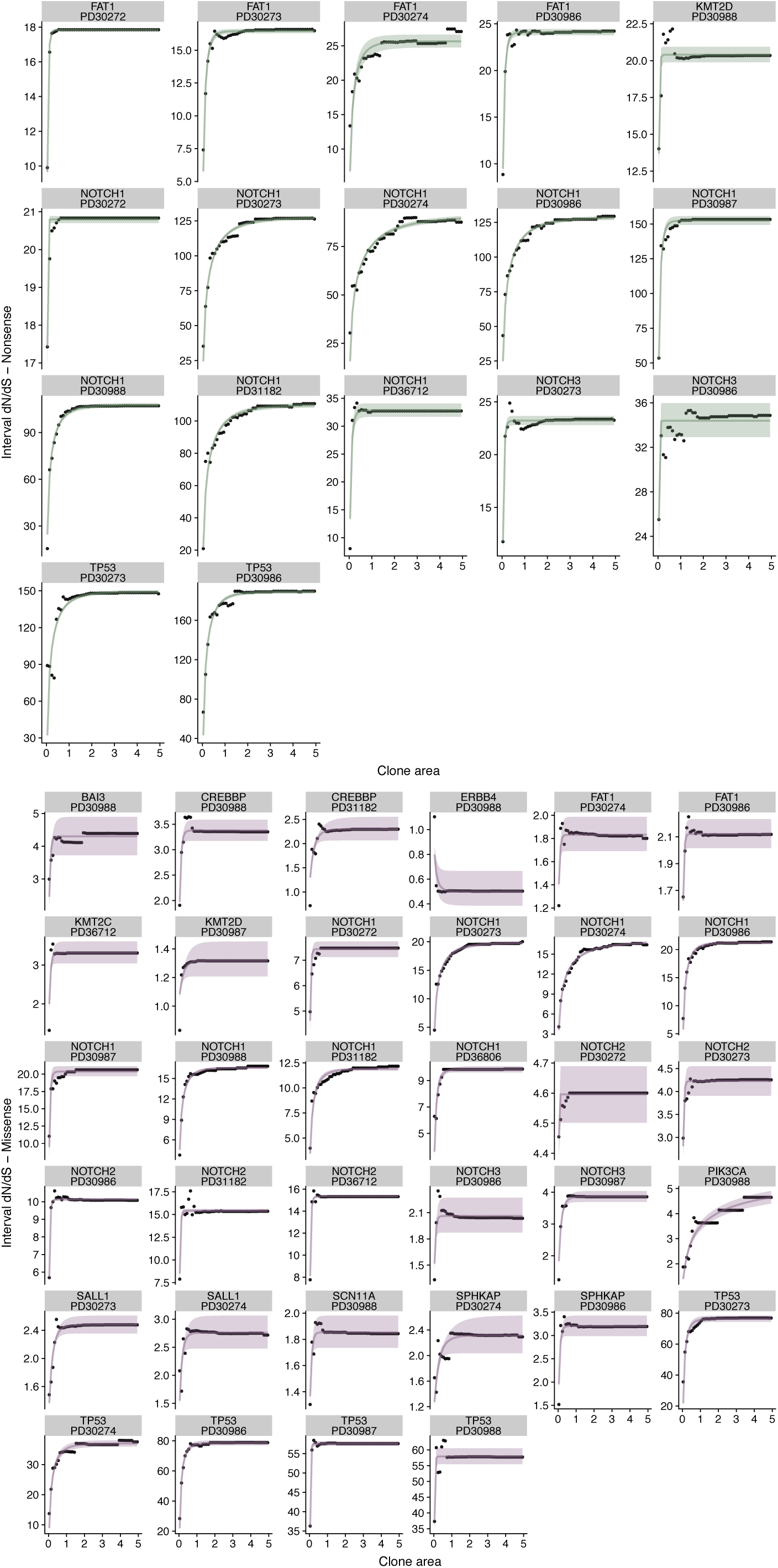
Individual fits for each gene in each patient in the oesophagus dataset. Points are data and lines are model fits. Analysis performed separately for nonsense, **a** and missense, **b**.

**Figure S5.**
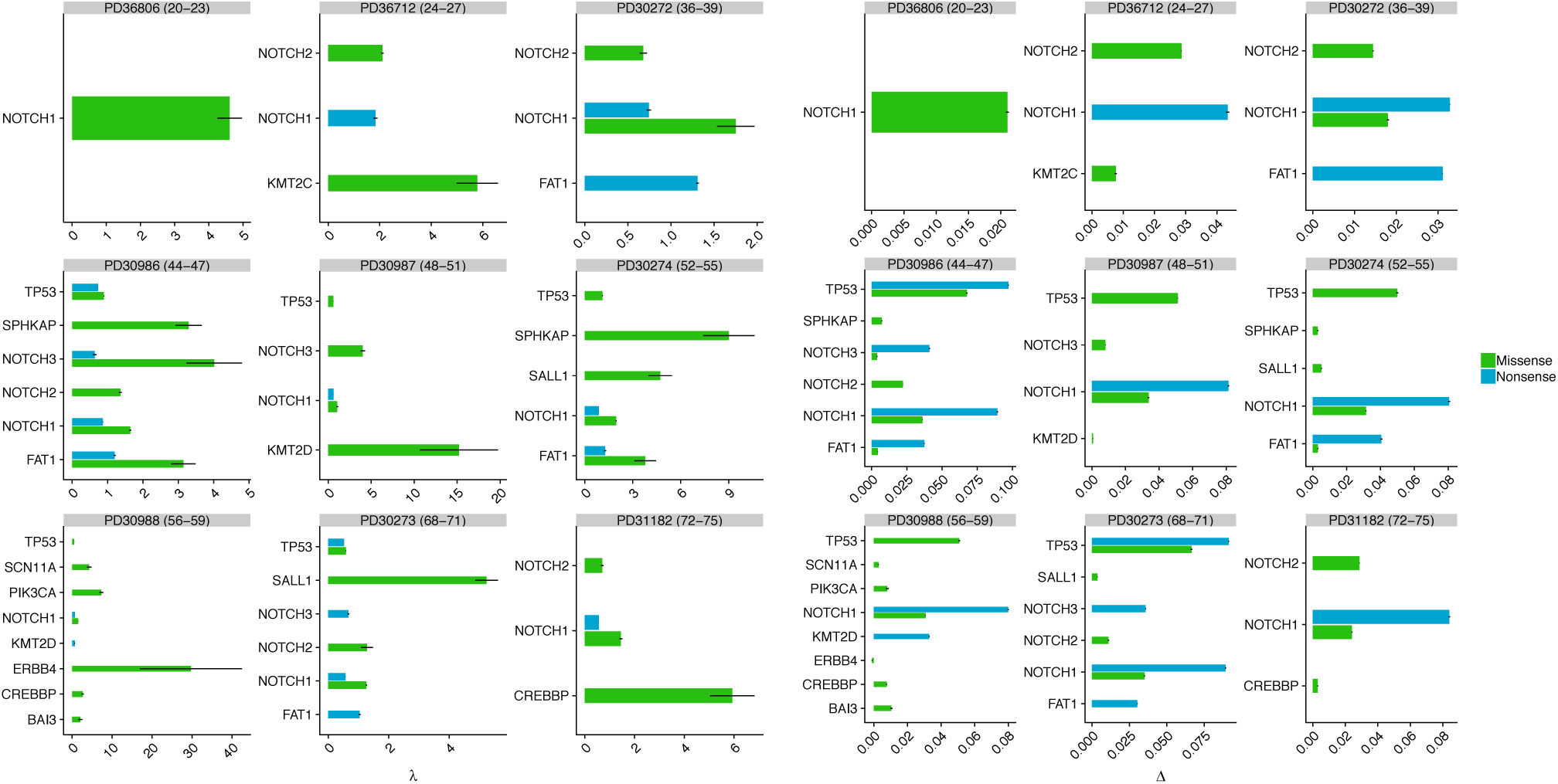
Inferred parameters for each gene in each patient in the oesophagus dataset where there were sufficient mutations to perform the analysis. Left hand plot shows inferred loss replacement rates *λ* and right hand plot inferred biases Δ.

**Figure S6.**
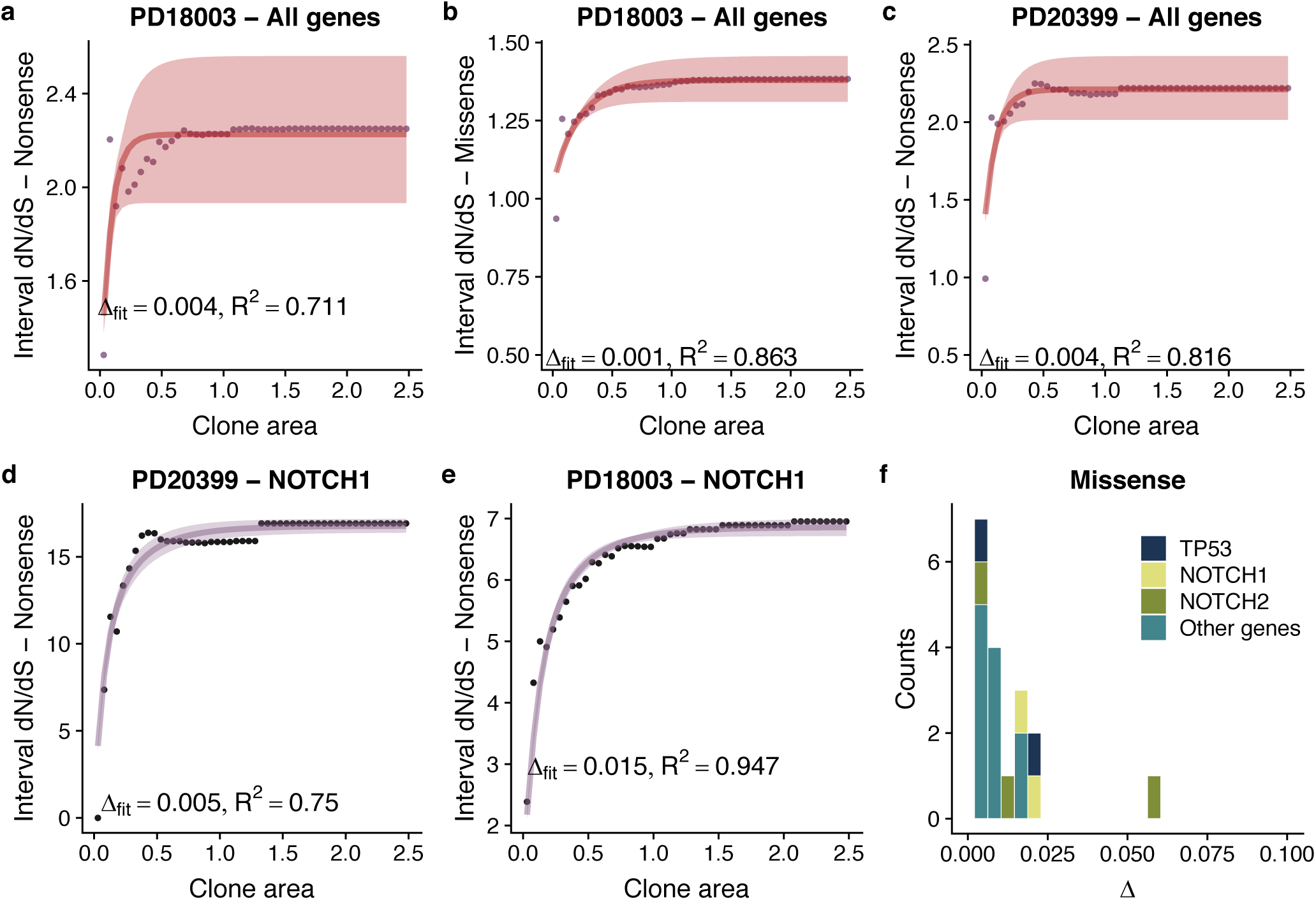
Model fits per patient and per gene per patient when there were sufficient mutations in the skin dataset. Points are data and lines are model fits, **a-e. f** shows the distributions of fitness effects for missense mutations across the cohort. There were insufficient nonsense mutations in the majority of genes to draw the equivalent plot for nonsense mutations.

**Figure S7.**
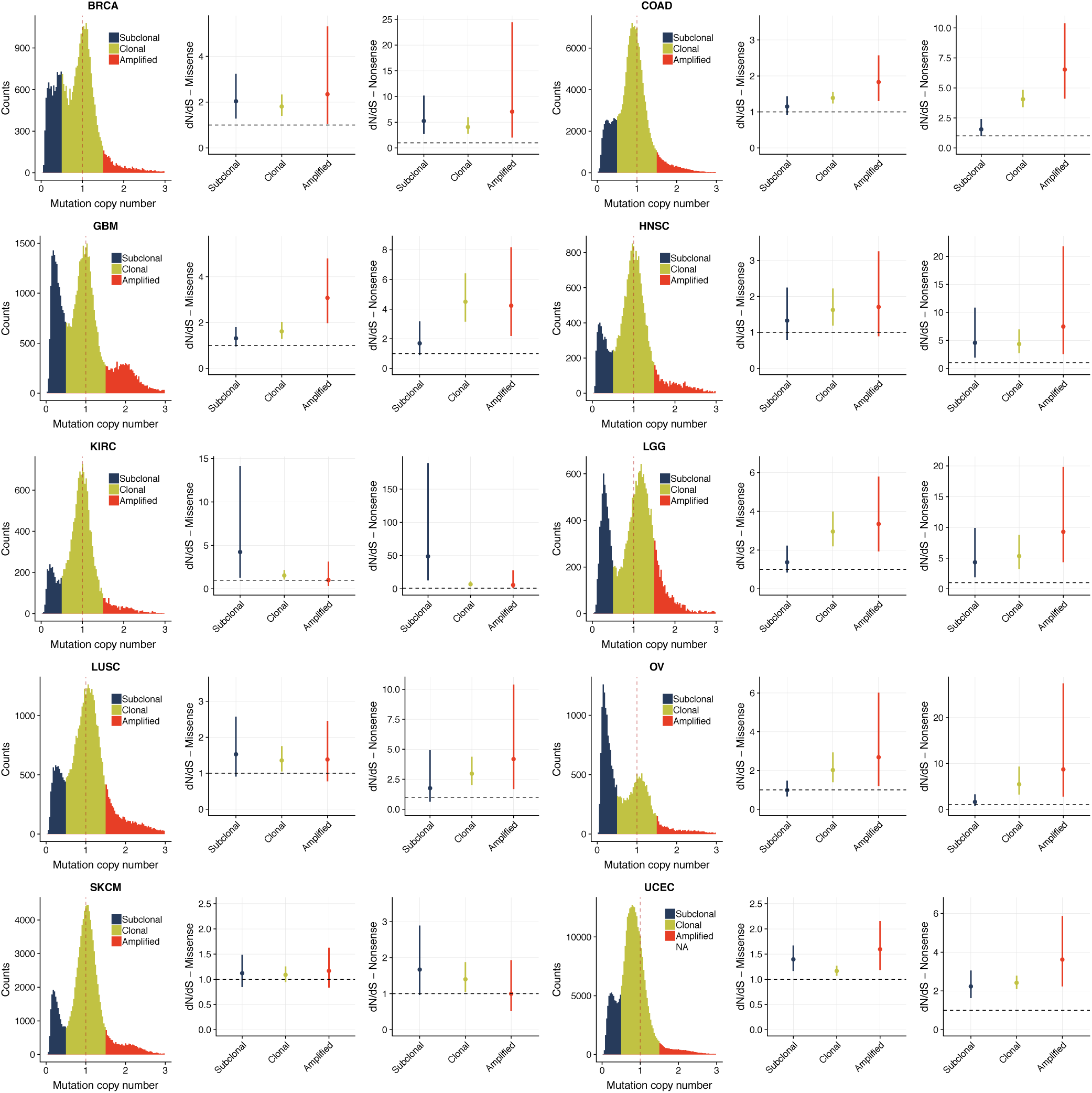
Mutation copy number histograms and dN/dS values for different cancer types with >100 samples (post filtering) in TCGA. Histograms and dN/dS plots coloured by mutation clonality.

**Figure S8.**
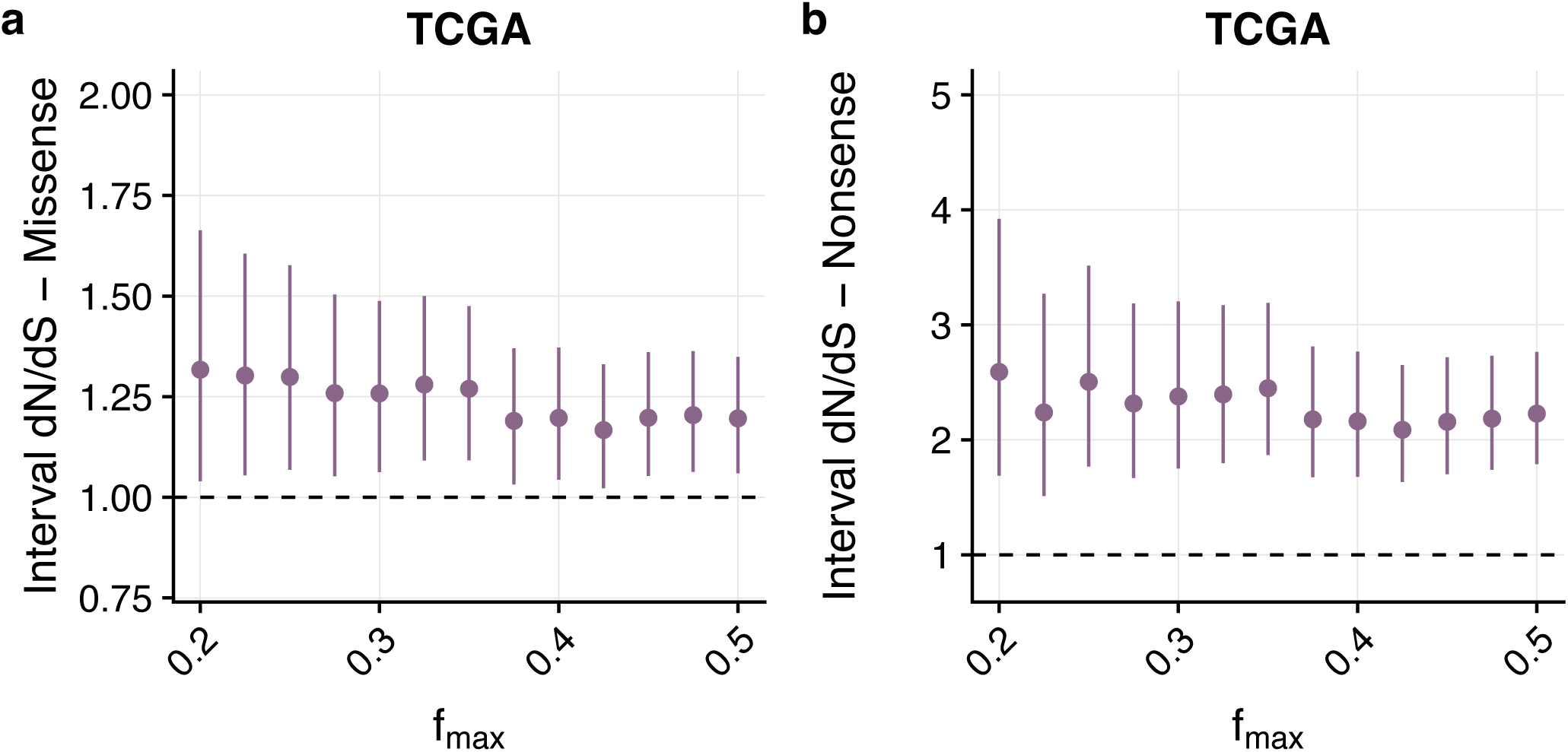
Interval dN/dS for 192 high confidence driver mutations. We observe no patterns that are predicted by our theoretical model.

**Figure S9.**
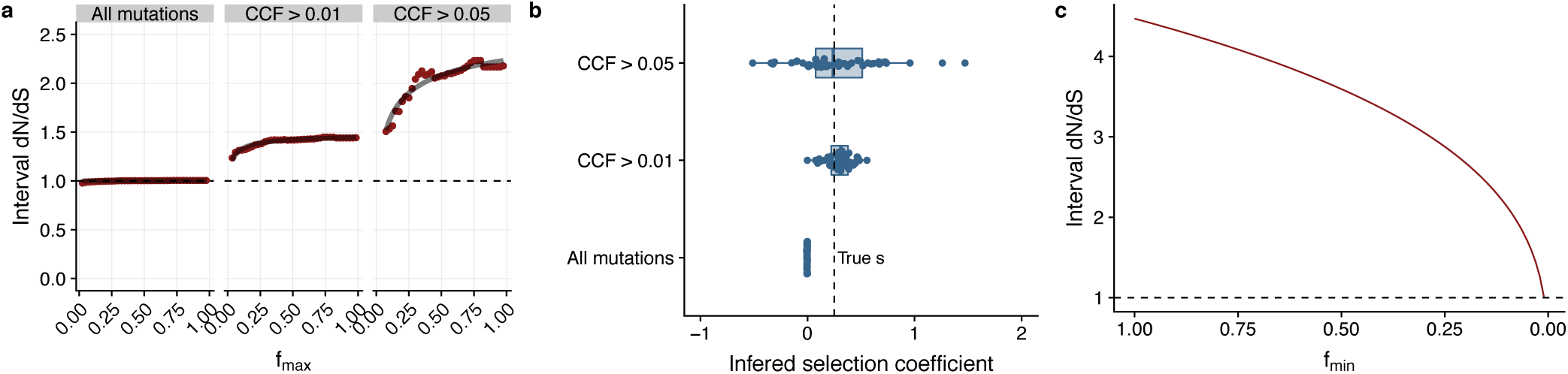
Generating a synthetic cohort with selection and using all mutations to infer dN/dS values shows that in this case dN/dS∼1, while if we restrict our attention to high frequency variants dN/dS>1, **a**. Inferred selection coefficients are accurate only when using high frequency variants, **b**. Using our theoretical interval model equation we see that fixing *f*_*min*_ = 1 and taking the limit *f*_*min*_ → 0 results in dN/dS = 1.

**Figure S10.**
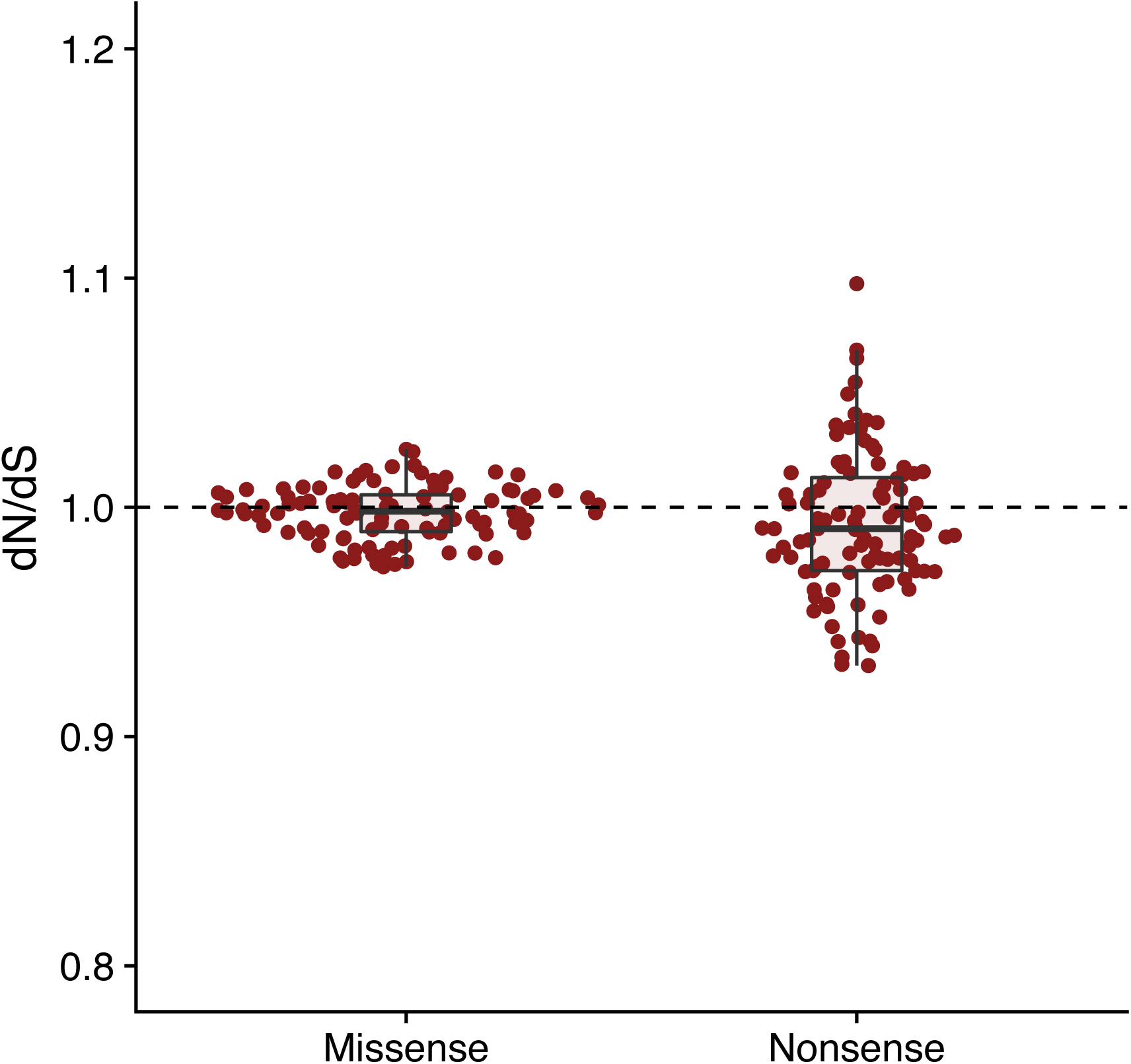
Corrected dN/dS values from 100 sets of 1000 randomly samples genes. Average dN/dS ∼ 1 as would be expected.

## References

1. Williams, M. J., Sottoriva, A. & Graham, T. Measuring Clonal Evolution in Cancer with Genomics. Annu. Rev. Genom. Hum. Genet. In Press

2. Eyre-Walker, A. & Keightley, P. D. The distribution of fitness effects of new mutations. Nat Rev Genet 8, 610–618 (2007).

3. Martincorena, I. et al. Universal Patterns of Selection in Cancer and Somatic Tissues. Cell 1–35 (2017). doi:10.1016/j.cell.2017.09.042

4. Vermeulen, L. et al. Defining stem cell dynamics in models of intestinal tumor initiation. Science 342, 995–998 (2013).

5. Rogers, Z. N. et al. Mapping the in vivo fitness landscape of lung adenocarcinoma tumor suppression in mice. Nature Genetics 50, 483–486 (2018).

6. Watson, C. J. et al. The Evolutionary Dynamics and Fitness Landscape of Clonal Haematopoiesis. BioRxiv 1–34 (2019). doi:10.1101/569566

7. Körber, V. et al. Evolutionary Trajectories of IDHWT Glioblastomas Reveal a Common Path of Early Tumorigenesis Instigated Years ahead of Initial Diagnosis. Cancer Cell 1–37 (2019). doi:10.1016/j.ccell.2019.02.007

8. Bailey, M. H. et al. Comprehensive Characterization of Cancer Driver Genes and Mutations. Cell 173, 371–385.e18 (2018).

9. Weghorn, D. & Sunyaev, S. Bayesian inference of negative and positive selection in human cancers. Nature Genetics 49, 1–8 (2017).

10. Zapata, L. et al. Negative selection in tumor genome evolution acts on essential cellular functions and the immunopeptidome. Genome Biology 19, 1–17 (2018).

11. Wu, C.-I., Wang, H.-Y., Ling, S. & Lu, X. The Ecology and Evolution of Cancer: The Ultra-Microevolutionary Process. Annu. Rev. Genet. 50, 347–369 (2016).

12. Greenman, C., Wooster, R., Futreal, P. A., Stratton, M. R. & Easton, D. F. Statistical analysis of pathogenicity of somatic mutations in cancer. Genetics 173, 2187–2198 (2006).

13. Yang, Z., Ro, S. & Rannala, B. Likelihood models of somatic mutation and codon substitution in cancer genes. Genetics 165, 695–705 (2003).

14. Martincorena, I. et al. Somatic mutant clones colonize the human esophagus with age. Science 57, eaau3879–14 (2018).

15. Lee-Six, H. et al. Population dynamics of normal human blood inferred from somatic mutations. Nature 14, 213–478 (2018).

16. Nielsen, R. & Yang, Z. Estimating the distribution of selection coefficients from phylogenetic data with applications to mitochondrial and viral DNA. Mol Biol Evol 20, 1231–1239 (2003).

17. McGranahan, N. & Swanton, C. Clonal Heterogeneity and Tumor Evolution: Past, Present, and the Future. Cell 168, 613–628 (2017).

18. Sottoriva, A. et al. A Big Bang model of human colorectal tumor growth. Nature Genetics 47, 209–216 (2015).

19. Kryazhimskiy, S. & Plotkin, J. B. The Population Genetics of dN/dS. PLOS Genet 4, e1000304–10 (2008).

20. Mugal, C. F., Wolf, J. B. W. & Kaj, I. Why Time Matters: Codon Evolution and the Temporal Dynamics of dN/dS. Mol Biol Evol 31, 212–231 (2013).

21. Simons, B. D. Deep sequencing as a probe of normal stem cell fate and preneoplasia in human epidermis. PNAS 113, 128–133 (2016).

22. Durrett, R. Population genetics of neutral mutations in exponentially growing cancer cell populations. The Annals of Applied Probability 23, 230–250 (2013).

23. Klein, A. M., Brash, D. E., Jones, P. H. & Simons, B. D. Stochastic fate of p53-mutant epidermal progenitor cells is tilted toward proliferation by UV B during preneoplasia. Proc. Natl. Acad. Sci. U.S.A. 107, 270–275 (2010).

24. Lopez-Garcia, C., Klein, A. M., Simons, B. D. & Winton, D. J. Intestinal stem cell replacement follows a pattern of neutral drift. Science 330, 822–825 (2010).

25. Vermeulen, L. et al. Defining stem cell dynamics in models of intestinal tumor initiation. Science 342, 995–998 (2013).

26. Williams, M. J., Werner, B., Barnes, C. P., Graham, T. A. & Sottoriva, A. Identification of neutral tumor evolution across cancer types. Nature Genetics 48, 238–244 (2016).

27. Williams, M. J. et al. Quantification of subclonal selection in cancer from bulk sequencing data. Nature Genetics 50, 895–903 (2018).

28. Bozic, I., Gerold, J. M. & Nowak, M. A. Quantifying Clonal and Subclonal Passenger Mutations in Cancer Evolution. PLoS Comput Biol 12, e1004731 (2016).

29. Ling, S. et al. Extremely high genetic diversity in a single tumor points to prevalence of non-Darwinian cell evolution. Proc. Natl. Acad. Sci. U.S.A. 112, E6496–505 (2015).

30. Simons, B. D. Reply to Martincorena et al.: Evidence for constrained positive selection of cancer mutations in normal skin is lacking. Proc. Natl. Acad. Sci. U.S.A. 113, E1130–E1131 (2016).

31. Martincorena, I., Jones, P. H. & Campbell, P. J. Constrained positive selection on cancer mutations in normal skin. Proc. Natl. Acad. Sci. U.S.A. 113, E1128–E1129 (2016).

32. Klein, A. M. & Simons, B. D. Universal patterns of stem cell fate in cycling adult tissues. Development 138, 3103–3111 (2011).

33. Doupé, D. P. et al. A single progenitor population switches behavior to maintain and repair esophageal epithelium. Science 337, 1091–1093 (2012).

34. Alcolea, M. P. et al. Differentiation imbalance in single oesophageal progenitor cells causes clonal immortalization and field change. Nature Cell Biology 16, 612–619 (2014).

35. Nicholson, M. D. & Antal, T. Universal Asymptotic Clone Size Distribution for General Population Growth. Bull. Math. Biol. 78, 2243–2276 (2016).

36. Ewens, W. J. Mathematical Population Genetics. 1–435 (2012).

37. Martincorena, I. et al. Tumor evolution. High burden and pervasive positive selection of somatic mutations in normal human skin. Science 348, 880–886 (2015).

38. Bielski, C. M. et al. Widespread Selection for Oncogenic Mutant Allele Imbalance in Cancer. Cancer Cell 1–24 (2018). doi:10.1016/j.ccell.2018.10.003

39. Zheng, Q. Progress of a half century in the study of the Luria–Delbrück distribution. Math Biosci (1999). doi:10.1016/S0025-5564(99)00045-0

40. Kessler, D. A. & Levine, H. Scaling Solution in the Large Population Limit of the General Asymmetric Stochastic Luria–Delbrück Evolution Process. J Stat Phys 158, 783–805 (2014).

41. Gerlinger, M. et al. Genomic architecture and evolution of clear cell renal cell carcinomas defined by multiregion sequencing. Nature Genetics 46, 225–233 (2014).

42. Lawrence, M. S. et al. Mutational heterogeneity in cancer and the search for new cancer-associated genes. Nature 499, 214–218 (2013).

43. Caravagna, G. et al. Detecting repeated cancer evolution from multiregion tumor sequencing data. Nat Methods 15, 1–13 (2018).

44. Korolev, K. S. et al. Selective sweeps in growing microbial colonies. Phys Biol 9, 026008 (2012).

45. Van den Eynden, J. & Larsson, E. Mutational Signatures Are Critical for Proper Estimation of Purifying Selection Pressures in Cancer Somatic Mutation Data When Using the dN/dS Metric. Front. Genet. 8, 415–9 (2017).

46. Cannataro, V. L., Gaffney, S. G. & Townsend, J. P. Effect Sizes of Somatic Mutations in Cancer. JNCI Journal of the National Cancer Institute 110, 1171–1177 (2018).

47. Temko, D., Tomlinson, I. P. M., Severini, S., Schuster-Böckler, B. & Graham, T. A. The effects of mutational processes and selection on driver mutations across cancer types. Nat Commun 9, 1857 (2018).

48. Heide, T. et al. Reply to ‘Neutral tumor evolution?’. Nature Genetics 48, 1–9 (2018).

49. Colaprico, A. et al. TCGAbiolinks: an R/Bioconductor package for integrative analysis of TCGA data. Nucleic Acids Research 44, e71–e71 (2016).

50. Martincorena, I. & Campbell, P. J. Somatic mutation in cancer and normal cells. Science 349, 1483–1489 (2015).

51. Bezanson, J., Edelman, A., Karpinski, S. & Shah, V. B. Julia: A Fresh Approach to Numerical Computing. SIAM Review (2017)

52. R Core Team, R: A Language and Environment for Statistical Computing, R Foundation for Statistical Computing, Vienna. 2018

